# SnapHiC-G: identifying long-range enhancer-promoter interactions from single-cell Hi-C data via a global background model

**DOI:** 10.1101/2023.03.01.530686

**Authors:** Weifang Liu, Wujuan Zhong, Paola Giusti-Rodríguez, Geoffery W. Wang, Ming Hu, Yun Li

## Abstract

Harnessing the power of single-cell genomics technologies, single-cell Hi-C (scHi-C) and its derived technologies provide powerful tools to measure spatial proximity between regulatory elements and their target genes in individual cells. Using a global background model, we propose SnapHiC-G, a computational method to identify long-range enhancer-promoter interactions from scHi-C data. We applied SnapHiC-G to scHi-C datasets generated from mouse embryonic stem cells and human brain cortical cells and demonstrated that SnapHiC-G achieved high sensitivity in identifying long-range enhancer-promoter interactions. Moreover, SnapHiC-G can identify putative target genes for non-coding GWAS variants, and the genetic heritability of neuropsychiatric diseases is enriched for single nucleotide polymorphisms (SNPs) within SnapHiC-G-identified interactions in a cell-type-specific manner. In sum, SnapHiC-G is a powerful tool for characterizing cell-type-specific enhancer-promoter interactions from complex tissues and can facilitate the discovery of chromatin interactions important for gene regulation in biologically relevant cell types.

## Introduction

Chromatin spatial organization plays an essential role in genome function and gene regulation^1–3^. Genome-wide chromatin conformation capture technologies like Hi-C have improved our understanding of the 3D genome structure and aided the discovery of chromatin features at various scales, such as chromosome territories, A/B compartments, topologically associating domains (TADs), and chromatin loops. In bulk Hi-C analysis, millions of cells are pooled together to generate a population-averaged chromatin contact map. However, the chromatin features obtained in a large population of cells are not representative of individual cells from a complex mixture^4,5^, since the dynamics of chromatin conformation in individual cells are masked. Although larger-scale organizational features of the genome are evolutionarily conserved and highly reproducible at the population level^6^, direct evidence from single-cell imaging- and sequencing-based technologies reveal considerable cell-to-cell variability in chromatin folding features at smaller units such as chromatin loops and interactions^7–13^, suggesting genome organization is remarkably more variable than expected^14^. In particular, chromatin interactions identified in bulk Hi-C data are typically present in only a relatively small proportion (10%∽30%) of the cell population at any given time^8,9^, indicating a high degree of cell-level heterogeneity, consistent with the stochastic gene expression widely observed in mammals^14^. Therefore, studying chromatin features in single cells can provide new insights into genome function and gene regulation.

Recent advances in single-cell genomics enable profiling of 3D genome structures in single cells at unprecedented throughput and resolution^7,11,12,15–21^. Single-cell Hi-C (scHi-C) adopts the bulk Hi-C protocol with an extra step to isolate or barcode single cells^15^. Like bulk Hi-C, the standard scHi-C data processing workflow includes read alignment, deduplication, filtering of contacts and cells, and binning to generate the final contact matrix ready for data analysis^15,16^. Specifically, chromatin contacts for every cell are represented by a symmetric matrix with entries representing contact frequencies between fixed-length genomic regions, referred to as bin pairs. Analysis of scHi-C data includes 3D architecture reconstruction and feature extraction (interactions, loops, TADs, A/B compartments) extraction for single cells^15,16^. Due to the extreme sparsity in scHi-C data (over 99% zeros in contact matrices at kilobase resolution) and limited capture efficiency, imputation of missing contacts is essential to enhance the signal-to-noise ratio and subsequently allow for the detection of chromatin features. The first scHi-C data imputation method is proposed in Zhou *et al*.^22^, which applies the random walk with restart (RWR) algorithm. An alternative method Higashi^23,24^, models both cells and bins as nodes and then uses a hypergraph neural network to learn edges between these nodes considering the similarities and variability between cells instead of imputing each cell separately. More recently, scVI-3D^25^ was developed to model interaction frequencies by a deep degenerative model with a zero-inflation component to account for sparsity in scHi-C data. scVI-3D can impute sparse scHi-C contacts and account for the impact of genomic distance, library size, and batch effects. Furthermore, with the imputed data, latent representations of scHi-C can be learned to reconstruct 3D genome structures like TADs^26^.

While computational methods tailored for scHi-C data are still under development, many approaches have been proposed to detect chromatin loops or statistically significant long-range chromatin interactions from bulk Hi-C data. They can be broadly grouped into two classes, namely global background-based methods and local background-based methods. Global background-based methods fit a global statistical model based on 1D genomic distance and assign *p*-values to each bin pair in the contact matrix by comparing the observed contact frequency to the expected contact frequency under the global background. In contrast, local background-based methods identify peaks in the Hi-C contact map that are local maxima with respect to their neighboring bin pairs. Our recent work SnapHiC and SnapHiC2^27,28^ perform data imputation by RWR first and then combine global and local background models to identify chromatin loops from single cells of the same cell type. While being the first and only existing method to detect chromatin loops from scHi-C data, most SnapHiC-identified chromatin loops are CTCF-anchored structural loops due to the local background model, while the sensitivity to identify enhancer-promoter interactions is relatively low. To the best of our knowledge, no method exists for explicitly identifying enhancer-promoter interactions from scHi-C data.

To fill this gap, we propose SnapHiC-G, a new computational approach based solely on a global background model to identify long-range enhancer-promoter interactions from scHi-C data. We applied SnapHiC-G to re-analyze scHi-C datasets generated from mouse embryonic stem cells (mESCs) and human brain cortical cells. We showed that SnapHiC-G outperformed SnapHiC and existing methods designed for bulk Hi-C data, achieving higher sensitivity with comparable precision in identifying long-range enhancer-promoter interactions.

## Results

### Overview of the SnapHiC-G algorithm

The SnapHiC-G algorithm consists of four components: (i) imputing chromatin contact probabilities in every single cell, (ii) distance-stratified normalization of imputed contact probabilities, (iii) filtering candidate bin pairs, and (iv) identifying statistically significant long-range enhancer-promoter interactions. Given single-cell contact matrices, SnapHiC-G first applies the RWR algorithm to generate imputed contact probability matrices for every single cell using a sliding window approach (**Methods: SnapHiC-G algorithm**). Next, the imputed contact probabilities are converted into distance stratified Z-scores to account for the dependence between contact probability and the 1D genomic distance between two bins. To identify enhancer-promoter interactions, SnapHiC-G filters bin pairs based on transcript start sites (TSS) and the available epigenetic annotations (e.g., H3K27ac ChIP-seq peaks or ATAC-seq peaks) to obtain a set of candidate bin pairs that span between gene promoters and *cis*-regulatory regions (**Methods: SnapHiC-G algorithm**). SnapHiC-G then defines enhancer-promoter interactions based on the global background by applying the one-sample t-test for each tested bin pair across all single cells belonging to the same cell type. Specifically, for each bin pair, the one-sided hypothesis test is conducted where the null hypothesis states that the bin pair’s average value of normalized contact probability across all single cells equals zero. The alternative hypothesis states that the average normalized contact probability is greater than zero. By default, bin pairs with FDR<0.1 and t-statistics>3 are identified as significant enhancer-promoter interactions (**Methods: SnapHiC-G algorithm**). In our analysis, all scHi-C data are binned into 10Kb resolution unless stated otherwise.

### Benchmarking with mouse embryonic stem cells (mESCs)

We applied SnapHiC-G, SnapHiC^27^, FitHiC2^29^, FastHiC^30^, HiC-ACT^31^, and HiC-DC+^32^ on single-cell Hi-C data generated from 742 mouse embryonic stem cells (mESCs)^18^, where the latter four are methods designed for bulk Hi-C data. We aggregated single-cell Hi-C data for all cells as a pseudo-bulk Hi-C sample as input for bulk Hi-C methods (**Methods: Identification of loops/interactions using other Hi-C methods**). To benchmark against other methods, a reference list of significant interactions was constructed using HiCCUPS-identified loops from deeply sequenced bulk Hi-C data^33^ and MAPS-identified significant interactions from H3K4me3 PLAC-seq^34^, cohesion HiChIP^35^, and H3K27ac HiChIP data^36^. We took the union of all reference interactions and kept only bin pairs with a genomic distance between 20Kb and 1Mb for evaluation. The same filtering step implemented in SnapHiC-G was applied to outputs from other methods to ensure a fair comparison.

When applying to the complete set of 742 mESCs, SnapHiC-G identified notably more significant enhancer-promoter interactions than other methods and reached a genome-wide power of 80%, recovering most of the 38,588 interactions in the reference list (**Figure 1A**; **Table 1**). Due to extreme data sparsity, bulk Hi-C methods missed most reference interactions without imputing single-cell contact probabilities. FitHiC2 performed the best among other methods, calling 3,476 interactions with a genome-wide power of 16%. FastHiC identified more interactions than FitHiC2 with a lower genome-wide power of 12%. As expected, SnapHiC identified few enhancer-promoter interactions due to the local background model. Although already tailored for sparse scHi-C data, SnapHiC performed similarly to HiC-ACT, a global background method. HiC-DC+ identified the fewest interactions among all methods with the lowest genome-wide power.

**Figure 1.**
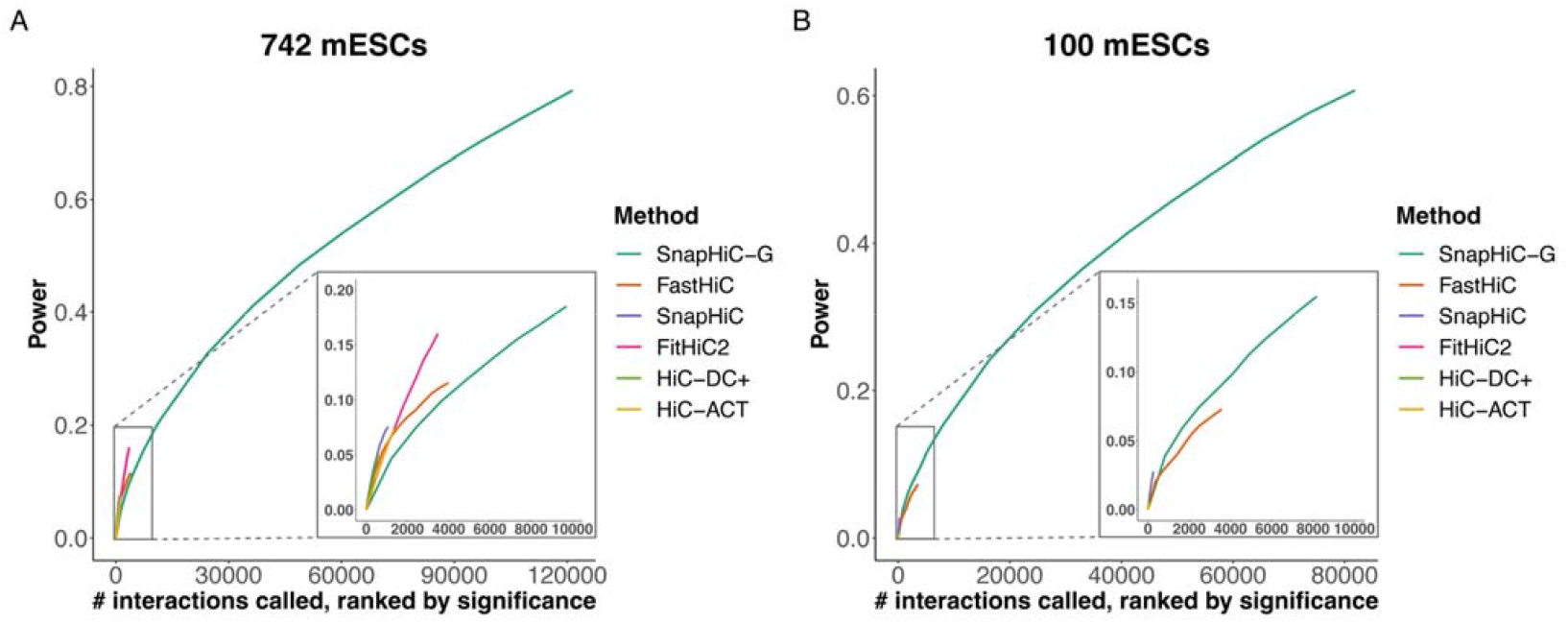
Power curves with (a) 742 mESCs and (b) 100 mESCs. Interactions were ranked by significance on the x-axis and power was evaluated with the corresponding number of top interactions. The lower right hand corner sub-figures are zoomed-in views of the top 10,000 interactions for each method.

**Table 1.** Genome-wide power and number of interactions called for mESCs and three brain cell types.

|  | 742 mESCs |  | 100 mESCs |  | 261 L2/3 neurons |  | 323 microglia |  | 1,038 oligodendrocytes |  |
| --- | --- | --- | --- | --- | --- | --- | --- | --- | --- | --- |
| Method | # interactions | Power | # interactions | Power | # interactions | Power | # interactions | Power | # interactions | Power |
| HiC-DC+ | 582 | 0.050 | 59 | 0.007 | 4,359 | 0.283 | 4,321 | 0.245 | 9,881 | 0.466 |
| FastHiC | 3,988 | 0.116 | 3,576 | 0.073 | 6,585 | 0.207 | 7,852 | 0.245 | 15,583 | 0.492 |
| HiC-ACT | 1,365 | 0.074 | 53 | 0.005 | 1,471 | 0.159 | 1,881 | 0.172 | 9,166 | 0.575 |
| FitHiC2 | 3,476 | 0.160 | 67 | 0.005 | 4,019 | 0.298 | 4,834 | 0.310 | 18,708 | 0.723 |
| SnapHiC | 1,070 | 0.076 | 265 | 0.028 | 2,918 | 0.203 | 1,766 | 0.147 | 3,605 | 0.242 |
| SnapHiC-G | 12,147 | 0.213 | 8,183 | 0.155 | 23,680 | 0.322 | 16,590 | 0.346 | 65,389 | 0.587 |

Since the number of input cells is critical in scHi-C data analysis, we assessed whether SnapHiC-G could retain its performance with fewer cells. Among all 742 mESCs, 100 mESCs were randomly selected, and the same performance evaluation of the six methods mentioned above was repeated. As shown in **Figure 1B** and **Table 1**, all methods had reduced power with 100 mESCs, while SnapHiC-G was least affected by the number of cells and showed more significant power gain over other methods compared with results from 742 cells. With only 100 mESCs, SnapHiC-G retained a genome-wide power of 61%, while all the other methods had genome-wide power below 10%. FastHiC still performed the best among others, with 3,576 significant interactions identified, comparable to the complete data, but the genome-wide power reduced almost by half to 7%. On the other hand, SnapHiC reached a 3% genome-wide power with 265 loops called; however, still more sensitive than other bulk Hi-C methods.

Due to the large number of significant interactions detected by SnapHiC-G, we evaluated the precision of identified interactions across the six methods with the same reference list. To control for the number of bin pairs compared, we ranked identified interactions in each method by their significance (i.e., *p*-value for HiC-ACT or FDR for FitHiC2, HiC-DC+, SnapHiC) or posterior probability (FastHiC) and calculated precision for the top 1,000, 2,000, 5,000, and 10,000 interactions. SnapHiC-G showed a comparable or better performance in terms of precision among the most significant interactions, even with a much larger number of interactions called (**Figure 2**). For example, with 742 mESCs, SnapHiC-G attained a precision of 0.93 for the top 1,000 interactions, which was comparable with HiC-DC+ (0.96) and FastHiC (0.96) and substantially higher than HiC-ACT (0.73), FitHiC2 (0.73), and SnapHiC (0.72). With 100 mESCs, SnapHiC-G had a precision of 0.60-0.76 for the top 1,000 to 10,000 interactions. We performed the same evaluation with three cell types from human brain cortical cells (oligodendrocytes, microglia, and L2/3 neurons) and had similar observations (**Supplementary notes; Figure S3; Figure S4**). Taken together, we have shown that SnapHiC-G had much higher sensitivity than other methods while maintaining precision among top enhancer-promoter interactions even with a small number of cells.

**Figure 2.**
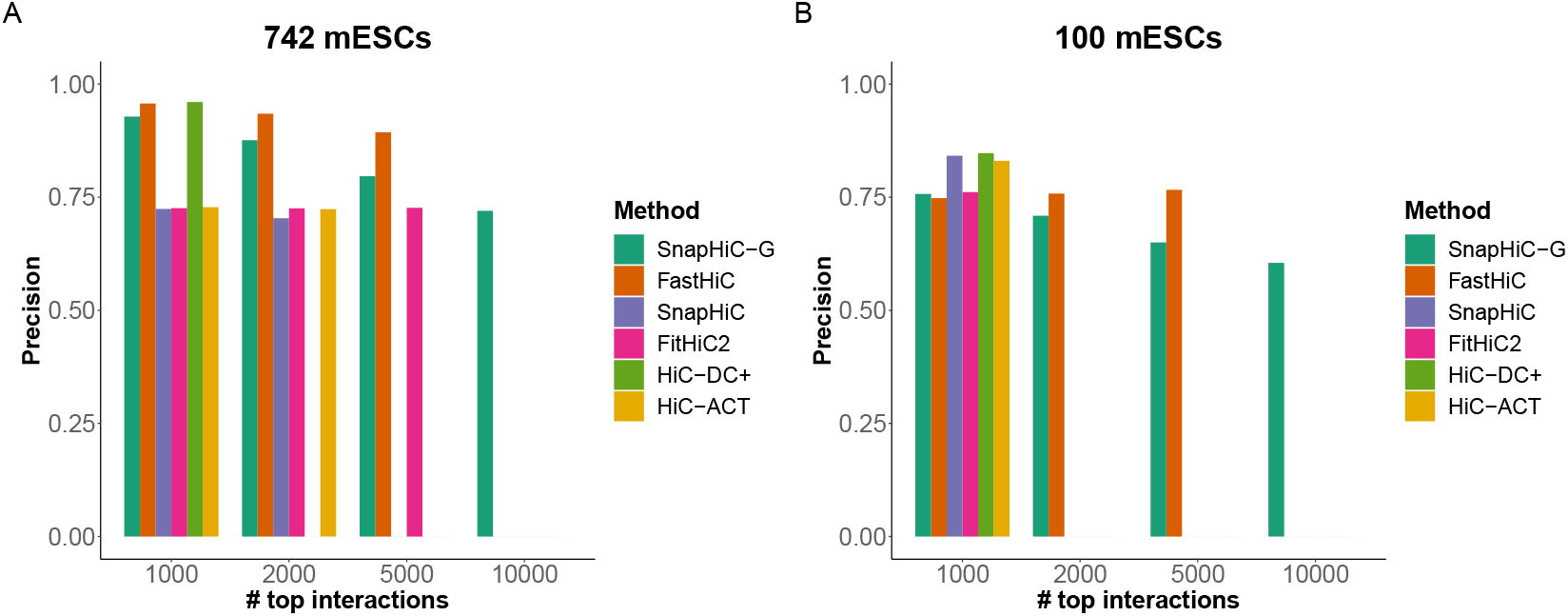
Precision bar plots with (a) 742 mESCs and (b) 100 mESCs. Results shown were for the top 1,000, 2,000, 5,000, and 10,000 interactions ranked by significance. Some bars are missing because the number exceeds the number of interactions called by that method.

### Enrichment of brain eQTL-TSS pairs in human cortical cells

To evaluate the functional characteristics of SnapHiC-G-identified enhancer-promoter interactions, we re-analyzed single-nucleus methyl-3C-seq (sn-m3c-seq) data from 2,869 human prefrontal cortical cells^37^. We collected eQTL data from various sources, including eQTL data from brain cell types released from Netherlands Brain Bank (NBB), the MS UK Tissue Bank (UKTB), and the Edinburgh Brain Bank (EBB)^38^, the CommonMind Consortium brain eQTL data^39^, and GTEx consortium v7 liver eQTL data^40^ as a control sample. For sn-m3c-seq data, cell types were classified based on DNA methylome as described in the original study^37^. We applied SnapHiC-G to four major cell types (astrocytes [n=338], microglia [n=323], oligodendrocytes [n=1038], and L2/3 neurons [n=261]) separately, where single cells from the same cell type were pooled (**Figure 3A; Table 1; Table S1**). We created control bin pairs to compute the enrichment of SnapHiC-G-identified enhancer-promoter interactions as eQTL-TSS of eGene pairs for each cell type. Specifically, we generated a pseudo bin pair for each significant enhancer-promoter interaction by retaining the bin with the promoter as the center but flipping the other bin to the opposite side of the center (**Methods: Enrichment of eQTL-TSS pairs**). We found that the odds for a bin pair to be an eQTL-TSS pair were significantly higher for significant interactions than pseudo bin pairs by overlapping with true eQTL-TSS bin pairs (**Figure 3A**). The wide confidence intervals (CI) of the odds ratios (OR) calculated from the Bryois *et al*. data^38^ were due to a smaller sample size to detect eQTLs compared with the CommonMind brain eQTL data (192 versus 467 brain samples, respectively). The two sets of brain eQTL data were highly consistent in terms of the order of enrichment for the four cell types, where oligodendrocytes showed the strongest enrichment, followed by astrocytes and microglia, and L2/3 neurons the weakest. As expected, SnapHiC-G-identified interactions were more enriched in brain eQTLs than in liver eQTLs. For example, the odds ratio for eQTLs from Bryois *et al*., CommonMind brain, and CommonMind liver in astrocytes were 2.35 (95% CI: 1.66 - 3.36), 1.95 (95% CI: 1.92 - 1.98), and 1.42 (95% CI: 1.29 - 1.55), respectively. These results validated the functional importance of SnapHiC-G-identified enhancer-promoter interactions in relevant cell types and tissues.

**Figure 3.**
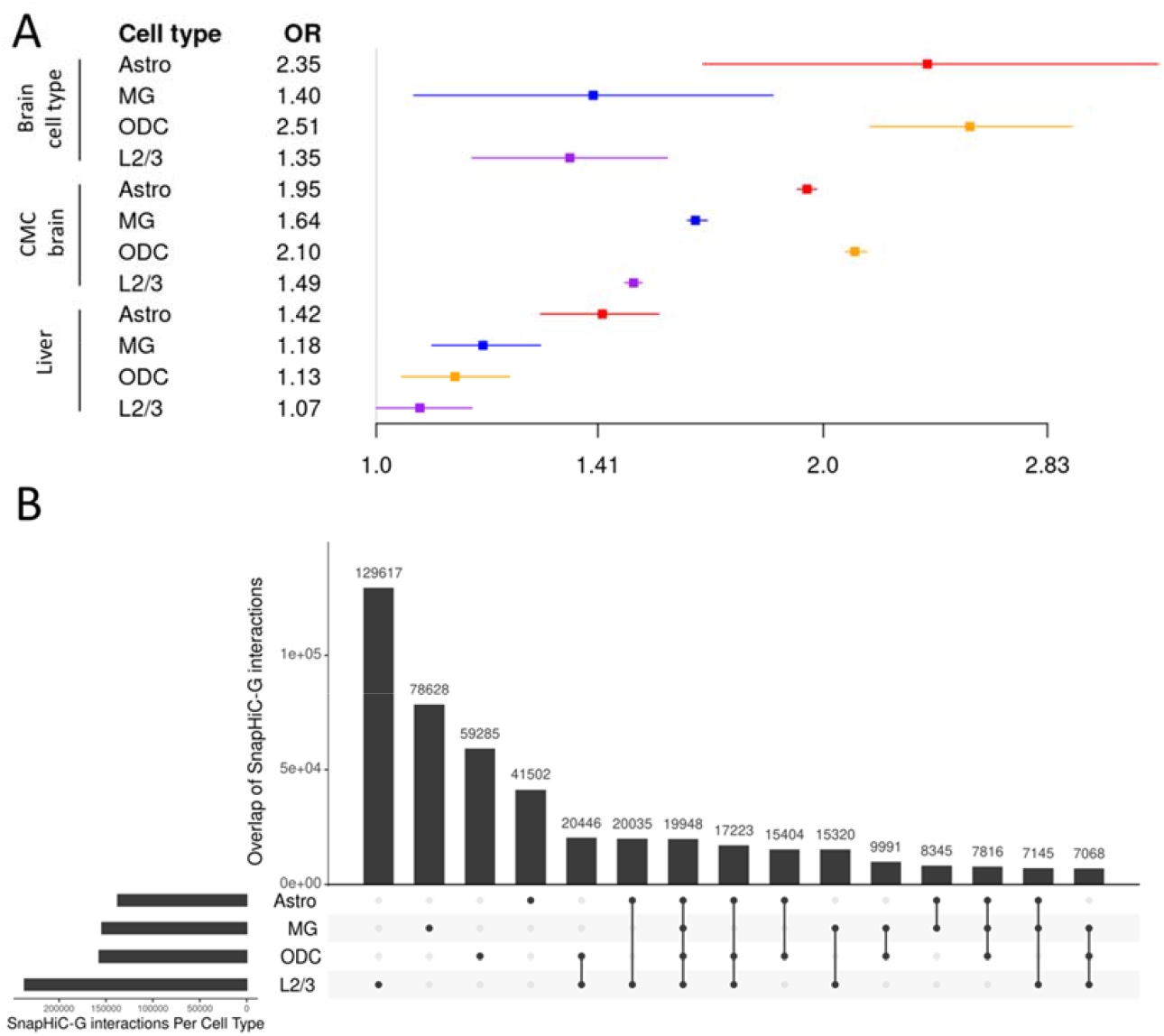
(A) Enrichment of SnapHiC-G-identified interactions for each brain cell type in CommonMind and liver eQTL-TSS pairs. The squares denote point estimates for odds ratios (ORs) and the error bars denote 95% confidence intervals for ORs. OR was defined as the odds of SnapHiC-G-identified interactions overlapping with eQTL-TSS bin pairs to the odds of pseudo bin pairs overlapping with eQTL-TSS bin pairs. (B) UpSet plot for SnapHiC-G-identified interactions. Number of exact overlapped interactions between four cell types: astrocytes (Astro), microglia (MG), oligodendrocytes (ODC), and L2/3 neurons (L2/3). The interactions were identified from 261 cells from each cell type. The UpSet figure is generated using the R package UpSetR.

### Gene expression patterns of cell-type-specific enhancer-promoter interactions

To further access the biological relevance of SnapHiC-G-identified enhancer-promoter interactions, we evaluated gene expression patterns in cell-type-specific interactions from the four brain cell types. Cell-type-specific interactions were defined as exclusively present in only one cell type, not including those shared by two or more cell types (**Figure 3B** and **Methods: Definition of overlapped enhancer-promoter interactions**). To avoid the impact of the number of cells in each cell type, we down-sampled astrocytes, microglia, and oligodendrocytes to 261 cells each to match the number of L2/3 neuron cells and identified cell-type-specific interactions with SnapHiC-G (**Methods: Down-sampling of scHi-C data**). Next, we selected genes with promoters overlapping with cell-type-specific interactions and extracted gene expression levels from RNA sequencing data from corresponding cell types^41^. As shown in **Figure 4A**, gene expression levels were higher for the genes from cell-type-specific enhancer-promoter interactions in the corresponding cell type. Moreover, after removing genes that overlapped with enhancer-promoter interactions detected by SnapHiC, a substantial number (above 65%) of the selected genes remained. We also observed increased expression of these genes in a cell-type-specific manner (**Figure 4B**). These results add another line of evidence that predicted enhancer-promoter interactions could provide valuable information in a cell-type-specific manner on top of the eQTL enrichment analysis.

**Figure 4.**
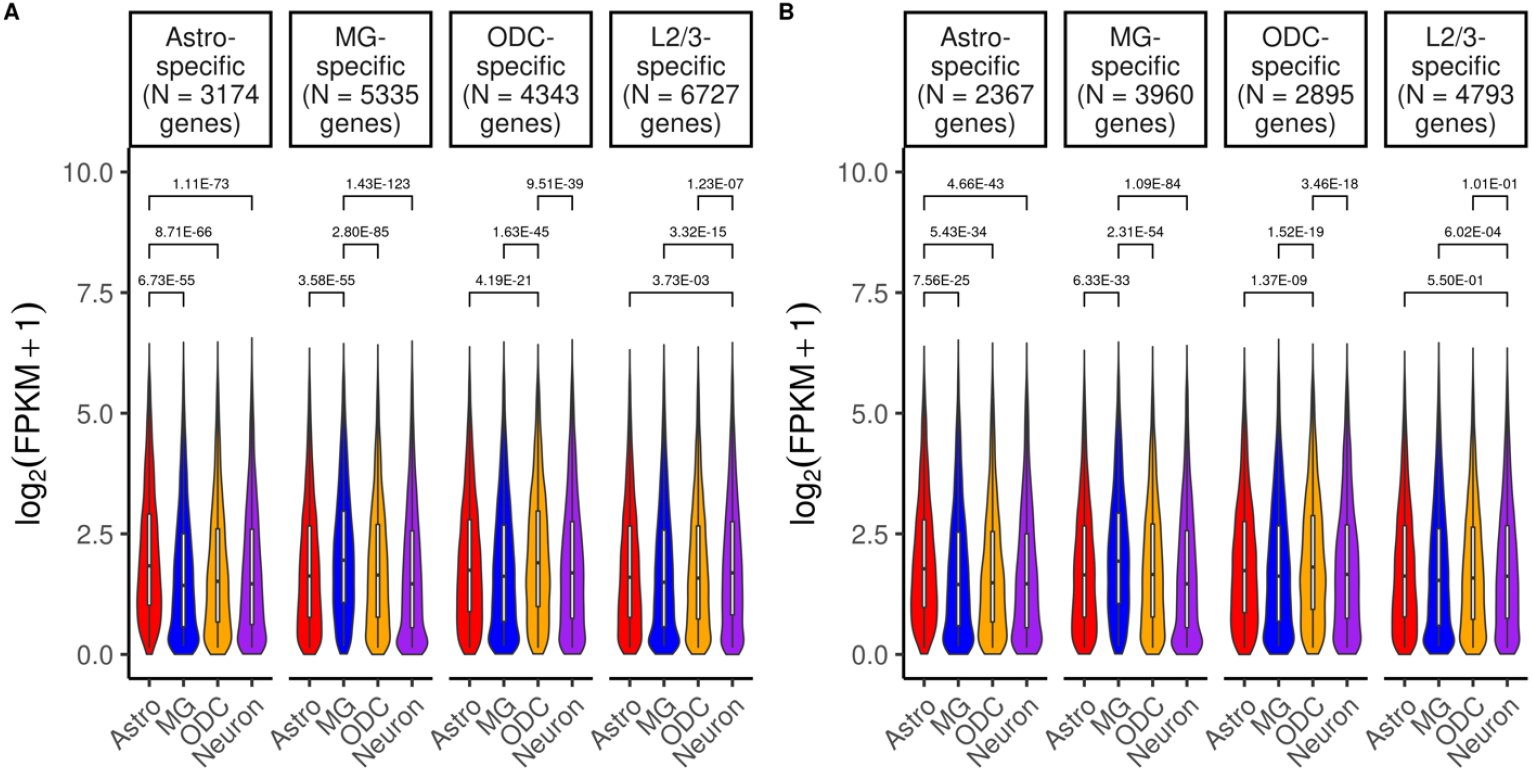
Cell-type-specific gene expression showing violin plots of RNA-seq expression levels (log2(FPKM+1) value) for selected genes. (A) Genes overlapping with cell-type-specific SnapHiC-G enhancer-promoter interactions. (B) Genes overlapping with cell-type-specific SnapHiC-G enhancer-promoter interactions, and those overlapping with enhancer-promoter interactions detected by SnapHiC were subsequently removed. P-values were calculated from paired Wilcoxon signed-rank tests. Gene expression outliers for each cell type were removed for visualization.

### Assigning GWAS variants to putative target genes

With over 90% of GWAS variants associated with human complex diseases and traits residing in non-coding regions yet enriched in *cis*-regulatory elements (e.g., promoters, enhancers, silencers, and insulators), enhancer-promoter interactions have the potential to prioritize disease-relevant genes for non-coding variants, particularly those in close spatial proximity that are far away in the 1D genomic distance with the promoter of their target genes. To assign putative target genes to non-coding GWAS variants based on predicted enhancer-promoter interactions in brain cell types, we collected the latest GWAS summary statistics for eight neurodevelopmental and neurodegenerative disorders: Alzheimer’s disease (AD)^42^, attention deficit hyperactivity disorder (ADHD)^43^, autism spectrum disorders (ASD)^44^, bipolar disorder (BP)^45^, schizophrenia (SCZ)^46^, Parkinson’s disease (PD)^47^, major depressive disorder (MDD)^48^, and neuroticism (NEU)^49^, and two complex traits: educational attainment (EDU)^50^ and intelligence quotient (IQ)^51^. We again focused on cell-type-specific enhancer-promoter interactions to predict target genes in a cell-type-specific manner using down-sampled sn-m3c-seq data. Specifically, SnapHiC-G identified 137,418, 154,261, 157,181, and 236,802 enhancer-promoter interactions in astrocytes, microglia, oligodendrocytes, and L2/3 neurons, respectively (**Table 2**). After excluding interactions shared among cell types, 14,440, 39,220, 23,853, and 65,819 cell-type-specific interactions were left correspondingly. To facilitate the interpretation of the GWAS variants, we focused on non-coding GWAS variants that reside in an active enhancer region in astrocytes, microglia, oligodendrocytes, or L2/3 neurons^52^. When matching GWAS variants and SnapHiC-G results, for each cell-type-specific enhancer-promoter interaction, we required that one bin contains GWAS variant(s) and the other bin overlaps with a gene’s TSS, and we annotated this gene as the putative target gene. Furthermore, we required that the corresponding gene is highly expressed (FPKM >1) in this cell type and lowly expressed (FPKM ≤1) in the other three cell types. We found 35, 82, 7, and 98 matched enhancer-promoter interactions (222 in total) for astrocytes, microglia, oligodendrocytes, and L2/3 neurons, respectively, and resolved over 600 SNP-disease associations (**Table 2**). Moreover, the average number of target genes for each variant was close to 1, much smaller than the number of nearby genes (+/- 1 Mb region), ranging from 25 to 83. For example, in astrocytes, the average number of target genes and nearby genes per variant were 1.1 and 38.3, respectively. These results showed that we could pinpoint target genes of non-coding variants in a cell-type-specific manner by integrating SnapHiC-G-identified enhancer-promoter interactions with GWAS results.

**Table 2.**
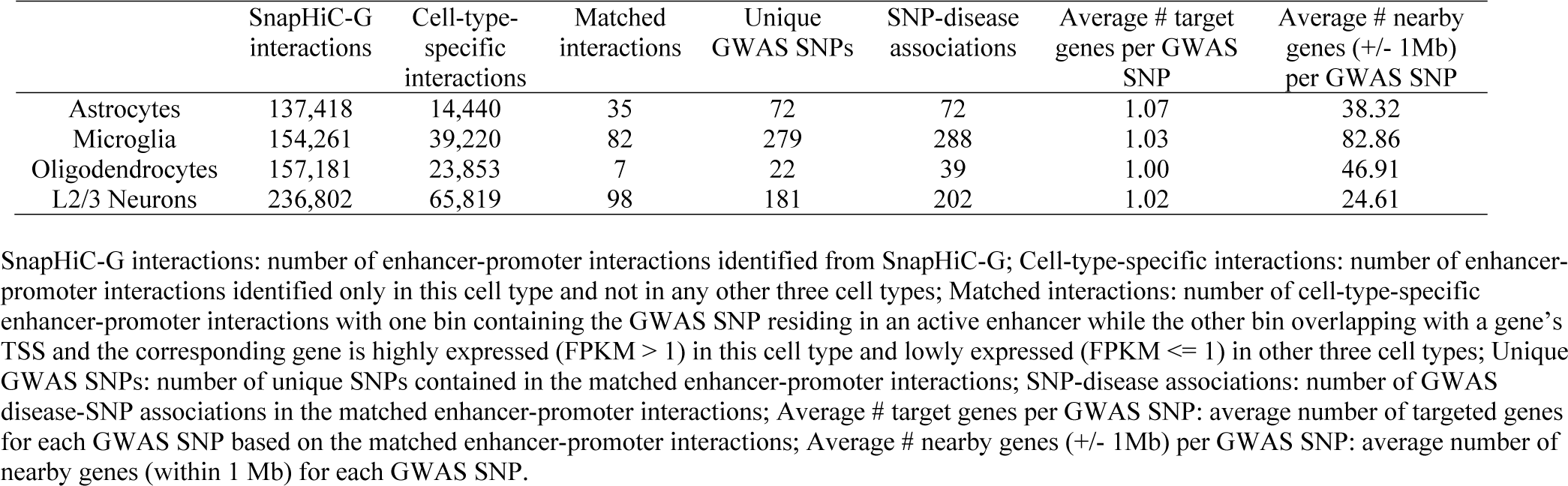
Summary of SnapHiC-G-identified enhancer-promoter interactions for each brain cell type.

|  | SnapHiC-G interactions | Cell-type-specific interactions | Matched interactions | Unique GWAS SNPs | SNP-disease associations | Average # target genes per GWAS SNP | Average # nearby genes (+/- 1Mb) per GWAS SNP |
| --- | --- | --- | --- | --- | --- | --- | --- |
| Astrocytes | 137,418 | 14,440 | 35 | 72 | 72 | 1.07 | 38.32 |
| Microglia | 154,261 | 39,220 | 82 | 279 | 288 | 1.03 | 82.86 |
| Oligodendrocytes | 157,181 | 23,853 | 7 | 22 | 39 | 1.00 | 46.91 |
| L2/3 Neurons | 236,802 | 65,819 | 98 | 181 | 202 | 1.02 | 24.61 |

SnapHiC-G interactions: number of enhancer-promoter interactions identified from SnapHiC-G; Cell-type-specific interactions: number of enhancer-promoter interactions identified only in this cell type and not in any other three cell types; Matched interactions: number of cell-type-specific enhancer-promoter interactions with one bin containing the GWAS SNP residing in an active enhancer while the other bin overlapping with a gene's TSS and the corresponding gene is highly expressed (FPKM > 1) in this cell type and lowly expressed (FPKM ≤ 1) in other three cell types; Unique GWAS SNPs: number of unique SNPs contained in the matched enhancer-promoter interactions; SNP-disease associations: number of GWAS disease-SNP associations in the matched enhancer-promoter interactions; Average # target genes per GWAS SNP: average number of targeted genes for each GWAS SNP based on the matched enhancer-promoter interactions; Average # nearby genes (+/- 1Mb) per GWAS SNP: average number of nearby genes (within 1 Mb) for each GWAS SNP.

### Examples of cell-type-specific enhancer-promoter interactions

From the 222 matched cell-type-specific enhancer-promoter interactions mentioned in the previous section, we were able to map a GWAS variant residing in an active enhancer with the promoter of a cell-type-specific gene (**Table 2**). Notably, most of these interactions were not identified by SnapHiC with the local background approach, as shown in **Figure 4B**. We demonstrate how SnapHiC-G-identified enhancer-promoter interactions can elucidate functional genes of GWAS loci in relevant cell types with a few examples.

The first example locates at a locus on chromosome 8, showing cell-type-specific interactions in L2/3 neurons and astrocytes **(Figure 5**). Specifically, a SCZ-associated GWAS SNP rs2565064 (chr8: 27,327,841) interacts with the promoter of *PNOC* in neurons, while another SCZ-associated SNP rs28541694 (chr8: 27,462,008) interacts with the promoter of *ZNF395* in astrocytes. In addition, ten AD-associated SNPs located in the chr8:27.4Mb-27.5Mb region are also connected to the promoter region of *ZNF395* in astrocytes, while none of the AD-associated SNPs interact with the *PNOC* promoter. These two genes also showed consistent cell-type-specific gene expression patterns, where *PNOC* was highly expressed in neurons (FPKM = 4.75 in neurons vs. ≤ 1 in the other three cell types) and *ZNF395* was highly expressed in astrocytes (FPKM = 2.26 in astrocytes vs. ≤ 1 in the other three cell types). Moreover, rs2565064 resides in a neuron-specific enhancer, and SNPs interacting with *ZNF395* reside in an astrocyte-specific enhancer. These results indicated that *PNOC* was the putative target gene for SCZ-associated SNP rs2565064 in neurons and *ZNF395* was the putative target gene for both SCZ and AD-associated SNPs in this 27.4Mb-27.5Mb region on chromosome 8. *PNOC* is primarily transcribed in the brain and spinal cord in the central nervous system^53^ and encodes the precursor for bioactive neuropeptides that influence a broad range of physiological roles, including memory, learning, and neuronal development, fear, anxiety, and sleep^54^. *PNOC* was also associated with PTSD, whose transcriptome significantly correlated with SCZ^55,56^. On the other hand, *ZNF395*, a gene involved in inflammation and cancer progression^57^, is upregulated in the SCZ network from the integrative network analysis^58^.

**Figure 5.**
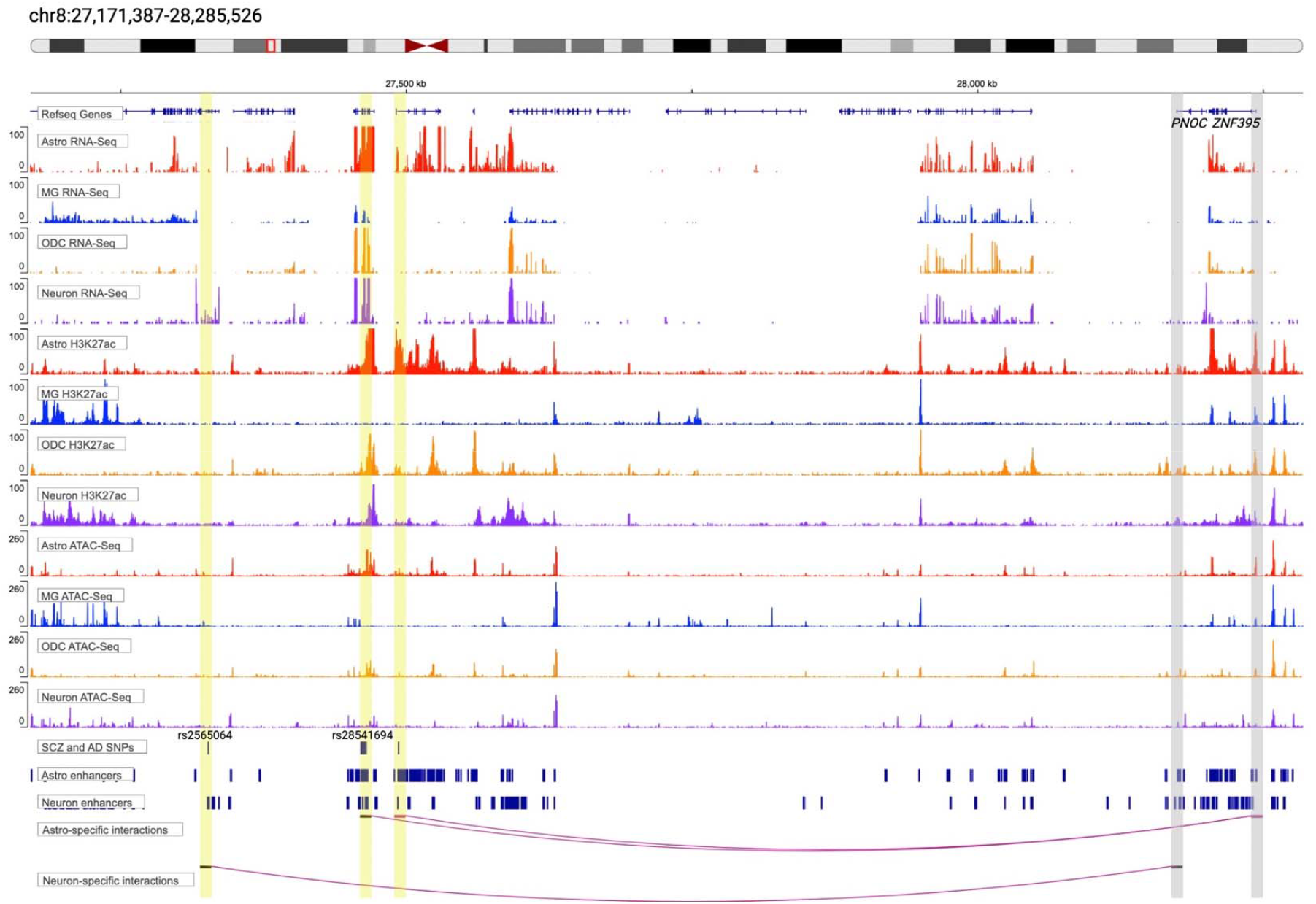
An illustrative example at the *PNOC-ZNF395* locus on chromosome 8 with neuron- and astrocyte-specific interactions. The top panel shows the gene track. The middle panels show RNA-seq, H3K27ac, and ATAC-seq tracks for the four brain cell types. The bottom panels show SCZ and AD GWAS SNPs, enhancer regions in astrocytes and neurons, and cell-type-specific interactions identified by SnapHiC-G but not SnapHiC. Astrocyte-specific interactions link the promoter region of *ZNF395* (highlighted in grey) with a SCZ-associated SNP rs28541694 and ten AD-associated SNPs located in the chr8:27.4Mb-27.5Mb locus (highlighted in yellow); a neuron-specific interaction links the promoter region of *PNOC* (highlighted in grey) with another SCZ-association SNP rs2565064 (highlighted in yellow). Both genes highlighted showed cell-type-specific gene expression in corresponding cell types. The anchors of these interactions also showed stronger H3K27ac ChIP-Seq and ATAC-seq signals in matched cell types.

Next, we focused on a microglia-specific interaction between the promoter of *ARPC1B*, which is highly expressed in microglia (FPKM = 7.12 in microglia vs. ≤ 1 in the other three cell types), and an AD-GWAS locus located in an active enhancer on chromosome 7 at 99.7 Mb (**Figure S1**). Our results showed that *ARPC1B* was the predicted target gene for this AD-associated GWAS variant rs1880949, consistent with prior findings that *ARPC1B* was active in microglia in AD patients but not in healthy controls^59^. At the same locus, SnapHiC-G detected another microglia-specific enhancer-promoter interaction between an active enhancer containing an EDU-associated GWAS variant rs10241492 (chr7: 99,994,813) and the *STAG3* gene’s promoter region. Moreover, rs10241492 was identified as an eQTL for *STAG3* from CommonMind in brain tissues^39^. In addition, *STAG3* was predicted to be the target gene for rs10241492 from a previous GWAS study of EDU^50^. These results together showed that the *STAG3* gene was potentially a target gene for rs10241492, specifically in microglia, consistent with findings in the original paper by Lee *et al*.^50^.

As a final example, **Figure S2** illustrates enhancer-promoter interactions identified specifically in microglia and neurons on chromosome 10. One microglia-specific enhancer-promoter interaction links a SCZ-associated GWAS locus in an active enhancer with *SFXN2*’s promoter. At the same time, *SFXN2* is highly expressed only in microglia (FPKM = 4.41 in microglia vs. ≤ 1 in the other three cell types). While a previous study suggested that *SFXN2* was a potential SCZ risk gene because of linkage or pleiotropic effects^60^, our results further indicated that *SFXN2* was the putative target gene for this particular SCZ-associated GWAS locus. Additionally, multiple neuron-specific enhancer-promoter interactions connect SCZ-associated GWAS loci to *INA* (FPKM = 46.21 in neurons vs. ≤ 1 in the other three cell types), suggesting this gene is a potential novel target gene for this locus with the evidence that corresponding GWAS variants (rs11191557, rs11191558, rs11191559, rs10883832, rs12413046) in this locus were also CommonMind eQTLs for *INA*.

Together, these examples showcase how SnapHiC-G-identified enhancer-promoter interactions can aid the interpretation of non-coding GWAS variants and reveal underlying mechanistic insights. Integrating with gene expression data and epigenetic annotations, SnapHiC-G was able to decipher the critical roles of non-coding variants in disease etiology in relevant cell types.

### Elucidating relevant cell types by heritability enrichment analysis

Next, we evaluated whether genetic heritability for complex diseases and traits was enriched for SNPs within the anchors of SnapHiC-G-identified interactions in specific cell types. Using the same set of GWAS summary statistics data together with two other complex traits: body mass index (BMI)^61^ and white blood cell count (WBC)^62^, we performed stratified linkage disequilibrium score regression (S-LDSC) analysis^63^. In brief, S-LDSC estimates the proportion of SNP heritability from predefined SNP-level functional annotations using GWAS summary statistics while accounting for linkage disequilibrium (LD) to identify functional categories enriched in SNP heritability and hence of functional relevance to the trait. In our case, the functional categories correspond to enhancer-promoter interactions from SnapHiC-G results in brain cell types, and our goal is to identify disease-relevant cell types for GWAS traits.

We first obtained SnapHiC-G-identified interactions from astrocytes, microglia, oligodendrocytes, and L2/3 neurons, all with 261 cells, to construct functional categories. As previously described, SnapHiC-G requires at least one bin overlapping with a TSS to ensure that the algorithm captures promoter-anchored interactions. Therefore, both bins may overlap with a TSS. However, to distinguish bins that overlap with a TSS and those that do not overlap with a TSS, we focused on the case where only one bin overlaps with a TSS. We focused on significant interactions in the four brain cell types to partition the genome. We defined SnapHiC-G anchor regions as the bins that overlap with TSS and SnapHiC-G target regions as the bins that do not overlap with TSS for each cell type. GWAS SNPs were annotated based on whether they fall into SnapHiC-G target regions for each cell type. Since the anchor regions all overlap with TSS, we evaluated whether SNPs located in SnapHiC-G target regions were enriched in SNP heritability in specific cell types.

**Figure 6** shows SNP heritability enrichment results for GWAS variants from the twelve traits analyzed, using SnapHiC-G target bins in four brain cell types as the functional categories. For example, AD SNP heritability was most strongly and significantly enriched in microglia target regions, which is consistent with the fact that the majority of AD GWAS risk loci are found close to genes highly expressed in microglia, and that microglia play a vital role in the pathogenesis of AD^64^. Notably, we observed two neuropsychiatric traits with high genetic correlation, namely bipolar disorder and schizophrenia, had enrichment scores and significance in the same order, where neuronal target regions were most strongly enriched, followed by astrocytes and oligodendrocytes. In contrast, enrichment in microglia was much lower. The strong enrichment in oligodendrocytes for Parkinson’s disease, although not the most significant, was consistent with the literature as well^65^. Results for white blood cell count showed that microglia were most strongly enriched, in agreement with the crucial roles that white blood cells have in the immune system.

**Figure 6.**
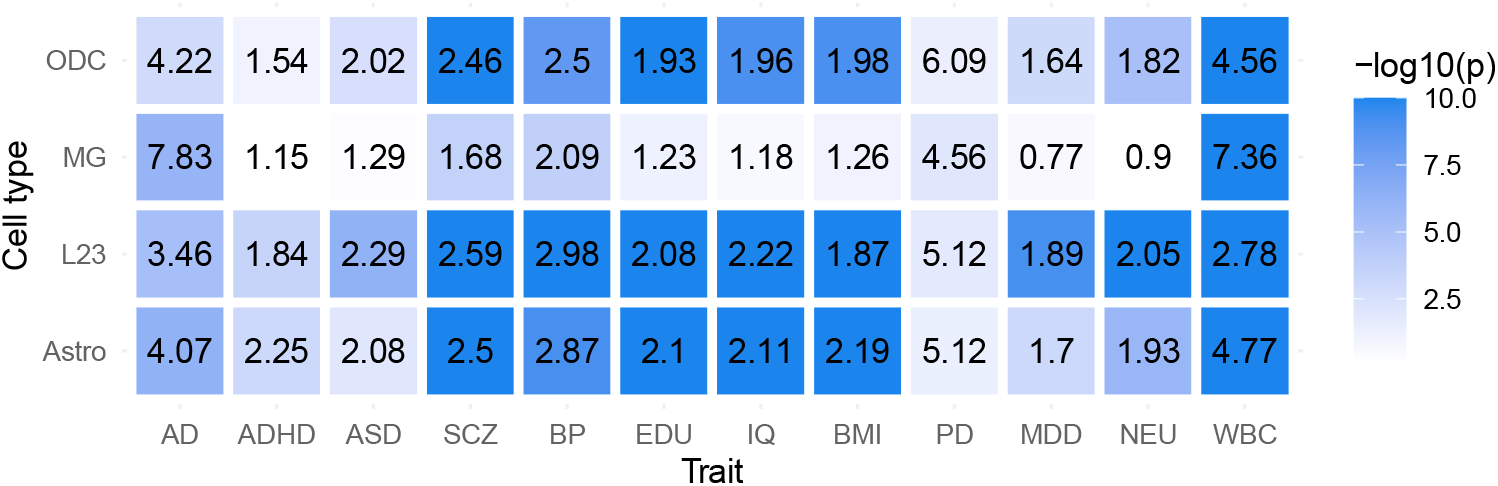
Heritability enrichment analysis results using LDSC. Cells were down-sampled to match the number of cells in L2/3 neurons. Numbers in the figure represent the enrichment score and colors represent the significance level in the -log10(p-value) scale. A higher enrichment score represents stronger enrichment in the corresponding cell type. Abbreviations: Alzheimer’s disease (AD), attention-deficit/hyperactivity disorder (ADHD), autism spectrum disorder (ASD), schizophrenia (SCZ), bipolar disorder (BP), educational attainment (EDU), intelligent quotient (IQ), body mass index (BMI), Parkinson’s disease (PD), major depressive disorder (MDD), neuroticism (NEU), white blood cell count (WBC).

## Conclusion and discussion

In this work, we developed SnapHiC-G, a computational pipeline to detect enhancer-promoter interactions from scHi-C data based on the global background model. To our knowledge, a method has yet to be proposed to address this task in scHi-C data. Our previous work, SnapHiC, the first computational pipeline to detect chromatin loops in scHi-C data, utilizes a combination of global and local background models to identify loop summits, therefore having limited power to detect enhancer-promoter interactions. Other existing global background methods, such as FitHiC2, HiC-DC+, FastHiC, and HiC-ACT, were designed for bulk Hi-C data and lacked the power to analyze sparse scHi-C contact data. To overcome the data sparsity challenge in scHi-C data, SnapHiC-G first applies the RWR algorithm to the observed raw contacts and then constructs a normalized contact probability matrix against the linear genomic distance. For each candidate bin pair, SnapHiC-G applies a one-sample t-test to test whether its average normalized contact frequency across single cells is significantly greater than zero (global background). Combined with epigenetic annotations such as enhancer and promoter marks, SnapHiC-G enables the profiling of cell-type-specific enhancer-promoter interactions by analyzing scHi-C data from multiple cell types.

Data from mESCs and human brain cortical cells showed that SnapHiC-G identified more significant interactions than existing global background-based methods designed for bulk Hi-C data with notably higher sensitivity. This was mainly due to the RWR imputation step, which significantly reduced the sparsity of the scHi-C data, especially when the number of cells was low. Aggregating a small number of single cells can result in very sparse pseudo-bulk Hi-C data, which may lead to poor performance of methods designed for bulk Hi-C. We evaluated the precision of top interactions identified by each method against a reference list of interactions and showed that SnapHiC-G did not suffer from a higher false discovery rate compared with other methods. The extra interactions called by SnapHiC-G may need further investigation, but we were not able to perform a systematic evaluation due to the lack of a gold-standard dataset.

Using GWAS summary statistics, we have also demonstrated the utility of enhancer-promoter interactions identified from SnapHiC-G in human brain cell types in prioritizing functionally relevant genes and cell types for complex human traits and diseases. SnapHiC-G can predict putative target genes for GWAS variants and help elucidate functionally relevant cell types of complex diseases.

For the human brain cell types, SnapHiC-G did not improve substantially over other methods compared with mESCs, especially when we look at the most significant interactions for oligodendrocytes (**Figure S3; Figure S4**). Several reasons can explain this. First, there was no gold-standard data for evaluating cell-type-specific enhancer-promoter interactions. The reference interaction lists inferred from H3K4me3 PLAC-seq data were suggestive rather than optimal, meaning that the reference might miss many cell-type-specific enhancer-promoter interactions. However, it was the best available data we could find. Second, SnapHiC-G-identified interactions had much lower FDRs than other bulk Hi-C methods. For identified oligodendrocyte interactions, the median of -log10(FDR) was 17.5 for SnapHiC-G, while the medians of other bulk Hi-C methods ranged from 1.7 to 7.5. The highly significant results of SnapHiC-G were due to both the RWR imputation and its global background nature; therefore, it can be hard to distinguish among identified interactions by ranking them by significance. Third, the number of cells for oligodendrocytes was relatively large (>1,000). After aggregating the single cells to construct bulk Hi-C data, it had a comparable sequencing depth to traditional bulk Hi-C data with ∽278 million intra-chromosomal contacts >20Kb. Consistent with SnapHiC, we observed a more significant power gain from SnapHiC-G when the number of cells was relatively small, which was consistent for other cell types.

SnapHiC-G performs the statistical test across all cells from the same cell type, which means that signals from the input cells are aggregated, and the identified enhancer-promoter interactions are still at the population level, similar to bulk Hi-C analysis. However, chromatin folding can be highly variable and dynamic even among cells of similar identities^18^, which is an exciting future direction. As more scHi-C data become available, multimodal data integration is another promising area of research to study the complexity and heterogeneity of chromatin interactions in single cells and can provide new insights into the regulatory mechanisms that underlie gene expression and cell differentiation^66^. scHi-C data also has great potential for predicting structural variations in cancer genomes, which is beyond the scope of this work^67^.

In our analysis, cell-type-specific epigenetic data considerably narrowed down candidate bin pairs. While such data aid the detection of cell-type-specific enhancer-promoter interactions, when they are not available, users can input only the TSS files to define the promoter regions and apply SnapHiC-G to identify promoter-interacting regions. With the rapid development of new technologies and more data available, scHi-C can trigger the study of fundamental questions about chromatin spatial organizations in individual cells during development, cancer cells, and different organs. Being able to identify cell-type-specific enhancer-promoter interactions from scHi-C data, SnapHiC-G results can be combined with the widely available GWAS results and epigenetic data, and it has a great potential to facilitate the discovery of regulatory chromatin interactions that are important for gene regulation in biologically relevant cell types.

## Methods

### SnapHiC-G algorithm

#### Step A. Imputation of contact probability using RWR with a sliding window approximation

We followed SnapHiC for imputing intra-chromosomal contact probability in every cell using the RWR algorithm following scHiCluster^22^. Each autosome was divided into consecutive 10 Kb bins, and each bin pair was converted to a binary representation with values 1 representing nonzero contact and 0 representing no contact observed. An unweighted and undirected graph modeled each chromosome by defining bins as the nodes and adjacent bin pairs or bin pairs with nonzero contact as edges. The RWR method was then used with a restart probability of 0.05 to estimate the likelihood of traveling between two nodes allowing for the imputation of contact probability between all intra-chromosomal bin pairs. The random-walk step captures information from global network structures, while the restart step captures information from local network structures.

Following SnapHiC2, we adopted a sliding window approach when imputing missing contacts to reduce computational costs. Specifically, instead of performing RWR over the entire chromosome, we divided the original contact matrix into partially overlapping matrices of size 2 Mb by 2 Mb along the diagonal line with overlapping areas of 1 Mb by 1 Mb. We then performed RWR for all 10 Kb bin pairs within each 2 Mb by 2 Mb submatrix along the diagonal to approximate contact probability. Only imputed contact probability in the middle rectangle areas was kept to avoid artifacts near corners.

#### Step B. Normalization of contact probability using 1D genomic distance

All bin pairs were stratified based on the genomic distance between two bins to account for the dependency between imputed contact probability and 1D genomic distance. Let *k* = 1,2, …, *K* be the index of *K* input cells. For a bin pair (*i, j*) in cell *k* with a genomic distance of *d*, let 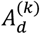 represent the strata including bin pairs in cell *k* with 1D genomic distance *d*. The normalized contact probability (*Z*-score) 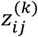 was calculated as 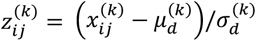, where 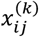 is the contact probability between bin *i* and bin *j* in cell *k*, 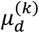 and 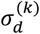 are mean and standard deviation of the contact probability of bin pairs in cell *k* within the strata 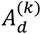.

#### Step C. Filtering the significant chromatin interactions

First, we define the “AND” bin pair as both sides overlapping with TSS and genes’ promoter regions, determined from the users’ input file or +/-500bp of TSS. Next, we define the “XOR” bin pair as only one side overlapping with TSS and genes’ promoter regions while the other side overlaps with enhancer regions. Significant chromatin interactions categorized as “AND” or “XOR” are the candidate SnapHiC-G enhancer-promoter interactions. Furthermore, candidate enhancer-promoter interactions with low mappability score (≤0.8) or overlapping with the ENCODE blacklist regions (mm10: http://mitra.stanford.edu/kundaje/akundaje/release/blacklists/mm10-mouse/mm10.blacklist.bed.gz; hg19: https://www.encodeproject.org/files/ENCFF001TDO/) were excluded. Our prior work was used to calculate each 10 Kb bin’s sequence mappability^68^.

#### Step D. Detecting enhancer-promoter interaction candidates

For each bin pair (*i, j*), we applied the one-sample *t*-test across all *K* cells to evaluate whether the contact probability was significantly higher than zero. We converted one sample *t*-test *p*-values into false discovery rates (FDRs), again stratified by 1D genomic distance. We defined a bin pair as a significant chromatin interaction if its (1) average of *z*-scores across all input cells > 0, (2) the proportion of cells with a *z*-score >1.96 (outlier cells) >10%, (3) FDR<10%, and (4) *t*-test statistic >3.

#### The computational cost of SnapHiC-G

Adopting a similar parallel computing strategy, SnapHiC-G is highly efficient regarding memory and computational time. We evaluated the computational cost of SnapHiC-G-specific steps (Step C and Step D) with both mouse and human scHi-C data under different settings and summarized the results in **Table S2**.

#### Processing scHi-C data

We used the same procedure as in SnapHiC to process the single-cell Hi-C data of mESCs and the human prefrontal cortex sn-m3C-seq data. For mESCs, we aligned the raw read pairs in fastq format to the mm10 genome, removed duplications, and then chose the top 742 cells (>150,000 contacts in each cell) for downstream analysis. For the human cortex, we used reference genome hg19 to process the data and then removed duplications. We chose the top 2,869 cells (>150,000 contacts in each cell) for downstream analysis and conducted cell type annotation as described in the original work.

#### Analyzing sn-m3C-seq data from the human cortex

We only analyzed data generated from the same cell type. For sn-m3C-seq data generated from complex tissue samples of heterogeneous cell types, we used cell type annotations reported by the original study^37^.

#### Down-sampling of scHi-C data

We randomly permuted 742 quality-controlled cells for the mESC scHi-C data to evaluate with down-sampled data. We then performed down-sampling by selecting the first 100 and 300 cells from the pool of 742 cells. As for the astrocytes (338 cells), microglia (323 cells), and oligodendrocytes (1,038 cells) from the human prefrontal cortex sn-m3C-seq data, we permuted the cells and selected the first 261 cells from each cell type to match the number of L2/3 neurons.

#### Identification of SnapHiC-G chromatin enhancer-promoter interactions

SnapHiC-G was applied to 742 mESCs (and down-sampled to 100 and 300 mESCs) scHi-C data and four cell types of the human prefrontal cortex sn-m3C-seq data to identify 10Kb bin chromatin enhancer-promoter interactions on autosomal chromosomes with 20Kb to 1Mb distance range. Bin pairs within 20 Kb were excluded from analyses.

#### Existing methods for bulk Hi-C data

Many methods, including FitHiC2, FastHiC, HiC-ACT, and HiC-DC+, have been developed for identifying long-range chromatin interactions from bulk Hi-C data. Specifically, FitHiC2^29,69^ is a spline regression-based approach to identify intra-chromosomal chromatin interactions. FastHiC^30^ is a Bayesian hidden Markov random field method that models the spatial dependency structure in high-resolution Hi-C data for detecting biologically meaningful chromatin interactions (i.e., peak calling). HiC-ACT^31^ is an aggregated Cauchy test-based approach that combines *p*-values from other methods (e.g., FitHiC2) without knowing the underlying spatial dependency structure. HiC-DC+^32^ is a negative binomial regression-based method based on genomic distance, GC content, and mappability features to predict chromatin interactions.

#### Identification of loops/interactions using other Hi-C methods

We identified chromatin loops/interactions with HiCCUPS, FastHiC, FitHiC2, HiC-ACT, and HiC-DC+ from aggregated contact matrix and SnapHiC from the single-cell contact matrix. To apply the methods developed for bulk Hi-C data, we generated pseudo bulk Hi-C data by aggregating 10Kb resolution scHi-C contact matrices across single cells (i.e., sum up the contacts)^27^. Then we applied FitHiC2, FastHiC, HiC-ACT, and HiC-DC+ to the pseudo bulk Hi-C data with lenient significance thresholds to consider the sparsity of scHi-C data. We used the following criteria for bulk Hi-C methods to detect significant interactions: FDR <10% from FitHiC2; posterior probability >0.9 from FastHiC; local neighborhood smoothed *p*-values <10^−6^ from HiC-ACT; FDR < 10% from HiC-DC+. For SnapHiC, we used FDR <10% and t-statistics >3 to select significant interactions. We further required interactions within the 20Kb-1Mb 1D genomic distance, with high mappability (>0.8) and no overlapping with ENCODE blacklist regions for standard quality control. To fairly compare with SnapHiC-G-identified interactions, we required interactions to be “AND” or “XOR” as an additional filtering criterion. Note that the number of interactions for SnapHiC was reduced compared with the original SnapHiC paper because of this additional filtering criterion.

#### Definition of overlapped enhancer-promoter interactions

We followed the same definition of overlapped loops as in SnapHiC. Let *d*_*im*_ denote the 1D genomic distance between the center of bin *i* and the center of bin *m*, and we define the distance between (*i, j*) and (*m, n*) as the maximum of *d*_*im*_ and *d*_*jn*_. For an enhancer-promoter interaction (*i, j*), if there exists an enhancer-promoter interaction (*m, n*) in set *S* such that the distance between (*i, j*) and (*m, n*) is within 20Kb, we define that the enhancer-promoter interaction (*i, j*) overlaps with the set *S*.

#### Definition of the reference interaction lists for evaluation

The reference interactions for mESCs consist of 10Kb interactions called from HiCCUPS with deeply sequenced bulk Hi-C data^33^ data, and MAPS-identified significant interactions from H3K4me3 PLAC-seq^34^, cohesion HiChIP^35^, and H3K27ac HiChIP data^36^. The reference interactions for oligodendrocytes, microglia, and neurons from the human prefrontal cortex were constructed using MAPS-identified H3K4me3 PLAC-seq data from corresponding cell types^52^, following SnapHiC. The same filtering steps as in SnapHiC-G and other methods were applied to the reference lists to ensure a fair evaluation.

#### Definition of cell-type-specific SnapHiC-G enhancer-promoter interactions

The cell-type-specific enhancer-promoter interactions were a subset of the SnapHiC-G enhancer-promoter interactions detected from down-sampled 261 cells in each of the four cell types: astrocytes, oligodendrocytes, microglia, and L2/3 excitatory neurons. Specifically, an enhancer-promoter interaction detected from a cell type was defined as a cell-type-specific SnapHiC-G enhancer-promoter interaction if none of the other three cell types’ identified enhancer-promoter interactions overlapped with it (**Methods: Definition of overlapped enhancer-promoter interactions**).

#### Enrichment of eQTL-TSS pairs

We evaluated whether SnapHiC-G-identified interactions are enriched in eQTL-TSS bin pairs in brain cell types, CommonMind brain eQTL-TSS bin pairs, and GTEx consortium liver eQTL-TSS bin pairs for all the cell types we considered. Specifically, for the eQTL-TSS bin pairs in brain cell types, we used eQTLs with Bonferroni corrected *p*-value < 0.05 within each cell type. Because the eQTL data did not have results in L2/3 neurons, we used eQTL results of excitatory neurons to compare with SnapHiC-G-identified bin pairs in L2/3 neurons. For each SnapHiC-G-identified interaction, we constructed a matched pseudo bin pair as a control: if only one bin contains the promoter, we kept the bin with the promoter as the center and flipped the other bin to be on the opposite side of the center but with the same distance from the center; if both bins contain promoters, we randomly selected one bin with probability 0.5 and kept it as the center, and then flipped the other bin to the other side of the center. Next, we removed the duplicates between controls and SnapHiC-G-identified bin pairs.

We constructed a two-by-two table for the union of SnapHiC-G-identified interactions and pseudo bin pairs, categorizing each bin pair by whether it is a SnapHiC-G-identified interaction or an eQTL-TSS bin pair. The *p*-value for independence between these two features was calculated using a two-sided Fisher’s exact test.

#### Gene expression analysis at cell type-specific enhancer-promoter interactions

The fragments per kilobase of transcript per million mapped reads (FPKM) values of each protein-coding gene in human astrocytes, neurons, microglia, and oligodendrocytes were acquired from Zhang *et al*.^41^. We used average FPKM values across biological replicates of the same cell type to quantify cell-type-specific gene expression levels.

#### Selection of GWAS SNPs associated with neuropsychiatric disorders and traits

First, we gathered significant (*p*-value <*5* × 10^−8^) GWAS SNPs for ten neuropsychiatric disorders and traits, including Alzheimer’s disease (AD)^42^, attention deficit hyperactivity disorder (ADHD)^43^, autism spectrum disorders (ASD)^44^, bipolar disorder (BP)^45^, schizophrenia (SCZ)^46^, Parkinson’s disease (PD)^47^, major depressive disorder (MDD)^48^, neuroticism (NEU)^49^, educational attainment (EDU)^50^ and intelligence quotient (IQ)^51^. Next, we overlapped these GWAS SNPs with active enhancers in four cell types (astrocytes, neurons, microglia, and oligodendrocytes), resulting in 9,764 SNP-trait associations (8,516 unique GWAS SNPs).

#### Web resources for figure generation

Figures 1, 2, 4, 6, S3, and S4 were created with the R package ggplot2 (https://CRAN.R-project.org/package=ggplot2). Figure 3A was created with the R package forestplot (https://CRAN.R-project.org/package=forestplot). Figure 3B was created with the R package UpSetR (https://CRAN.R-project.org/package=UpSetR). Figures 5, S1, and S2 were created with IGV (https://igv.org/app). Figures 1, 5, S1, S2, and S3 were arranged by BioRender (https://biorender.com).

## Data availability

Single-cell Hi-C data from mESCs were downloaded from https://www.ncbi.nlm.nih.gov/geo/query/acc.cgi?acc=GSE94489. sn-m3C-seq data from human prefrontal cortical cells were downloaded from https://www.ncbi.nlm.nih.gov/geo/query/acc.cgi?acc=GSE130711. The ATAC-seq and H3K27ac ChIP–seq data from human astrocytes, oligodendrocytes, microglia, and neurons were downloaded from dbGap https://www.ncbi.nlm.nih.gov/projects/gap/cgi-bin/study.cgi?study_id=phs001373.v2.p2. The RNA-seq data from human astrocytes, oligodendrocytes, microglia, and neurons were downloaded from https://www.ncbi.nlm.nih.gov/geo/query/acc.cgi?acc=GSE73721.

## Code availability

Our proposed SnapHiC-G method is implemented in the SnapHiC-G Python package, freely available on GitHub: https://github.com/wjzhong/SnapHiC-G.

## Acknowledgments

This study was funded by the NIH grants R35HG011922 (to M.H.) and U01DA052713 (to Y.L.). Y.L. was also partially supported by R01MH125236, and W.L. was supported by R01NR019245. We also thank assistance from the UNC Intellectual and Developmental Disabilities Research Center (NICHD; P50 HD103573).

## Author contributions

This study was conceived and designed by M.H. and Y.L. Data analysis was conducted by W.L. and W.Z. The SnapHiC-G software package was developed by W.Z. and W.L. The manuscript was written by W.L. and W.Z. with input from M.H., Y.L., P.G.R., and G.W.W.

## Competing interests

Author W.Z. was employed by Merck Sharp & Dohme LLC, a subsidiary of Merck & Co., Inc., Rahway, NJ, USA. The remaining authors declare that the research was conducted in the absence of any commercial or financial relationships that could be construed as a potential conflict of interest.

