## Supplementary materials for "SnapHiC-G: identifying long-range enhancer-promoter interactions from single-cell Hi-C data via a global background model"

### Supplementary notes

#### *Benchmarking with human brain cortical cells*

To evaluate the performance of SnapHiC-G in human brain cortical cells, we tested SnapHiC, FitHiC2, FastHiC, HiC-ACT, and HiC-DC+ along with SnapHiC-G on three cell types (oligodendrocytes, microglia, and L2/3 neurons) from 2,869 human brain cortical cells, each with more than 150,000 contacts, from the Lee *et al.* study[1], where bulk H3K4me3 PLAC-Seq data was available for reference (oligodendrocytes, microglia, and neurons)[2]. We applied each method for the three brain cell types separately and pooled single cells from the same cell type as the pseudo-bulk Hi-C data for bulk Hi-C methods. As shown in **Figures S3-S4**, SnapHiC-G detected the largest number of significant interactions and achieved higher sensitivity than alternative methods for the analyses of all three brain cell types, where the power gain was more substantial for L2/3 neurons and microglia. Similar to mESCs, there was a large difference between the number of significant interactions identified from different methods based on comparable significance thresholds (**Methods: Identification of loops/interactions using other Hi-C methods**). For example, with 261 L2/3 neurons, the number of significant interactions ranged from 1,471 for HiC-ACT to 6,585 for FastHiC, while SnapHiC-G identified more than 20,000 significant interactions (**Table 1**).

#### *Difference between SnapHiC and SnapHiC-G*

For each bin pair, SnapHiC-G applies the one-sample t-test to test whether its average normalized contact frequency across single cells is significantly greater than zero (global background), while SnapHiC requires the average value to be positive and larger than the mean value of its surrounding bin pairs (local background). As a result, SnapHiC-G called more interactions than SnapHiC, with a significantly higher sensitivity for detecting enhancer-promoter interactions with the same FDR cutoff.

The reduced sensitivity of SnapHiC can be explained by three reasons. First, gene promoters can form wide-spread interaction clusters with multiple narrow typical enhancers or broad super-enhancers, and enhancer-promoter interactions may not be located exactly at the summit of such clusters. In addition, CTCF-anchored loops can bring enhancers to the proximity of distal promoters to facilitate enhancer-promoter interaction, as suggested by the loop extrusion model[3,4]. Finally, enhancer-promoter interactions can be formed by the phase separation mechanism, which is independent of CTCF binding. In all these three scenarios, applying a local background model and selecting interaction summits will miss many of the enhancer-promoter interactions. Another key distinction between SnapHiC-G and SnapHiC is using cell-type-specific epigenetic data to narrow down candidate bin pairs. While such data greatly aid the detection of cell-type-specific enhancer-promoter interactions, when they are not available, users can input only the TSS files to define the promoter regions and apply SnapHiC-G to identify promoter-interacting regions.

**Supplementary figures and tables**

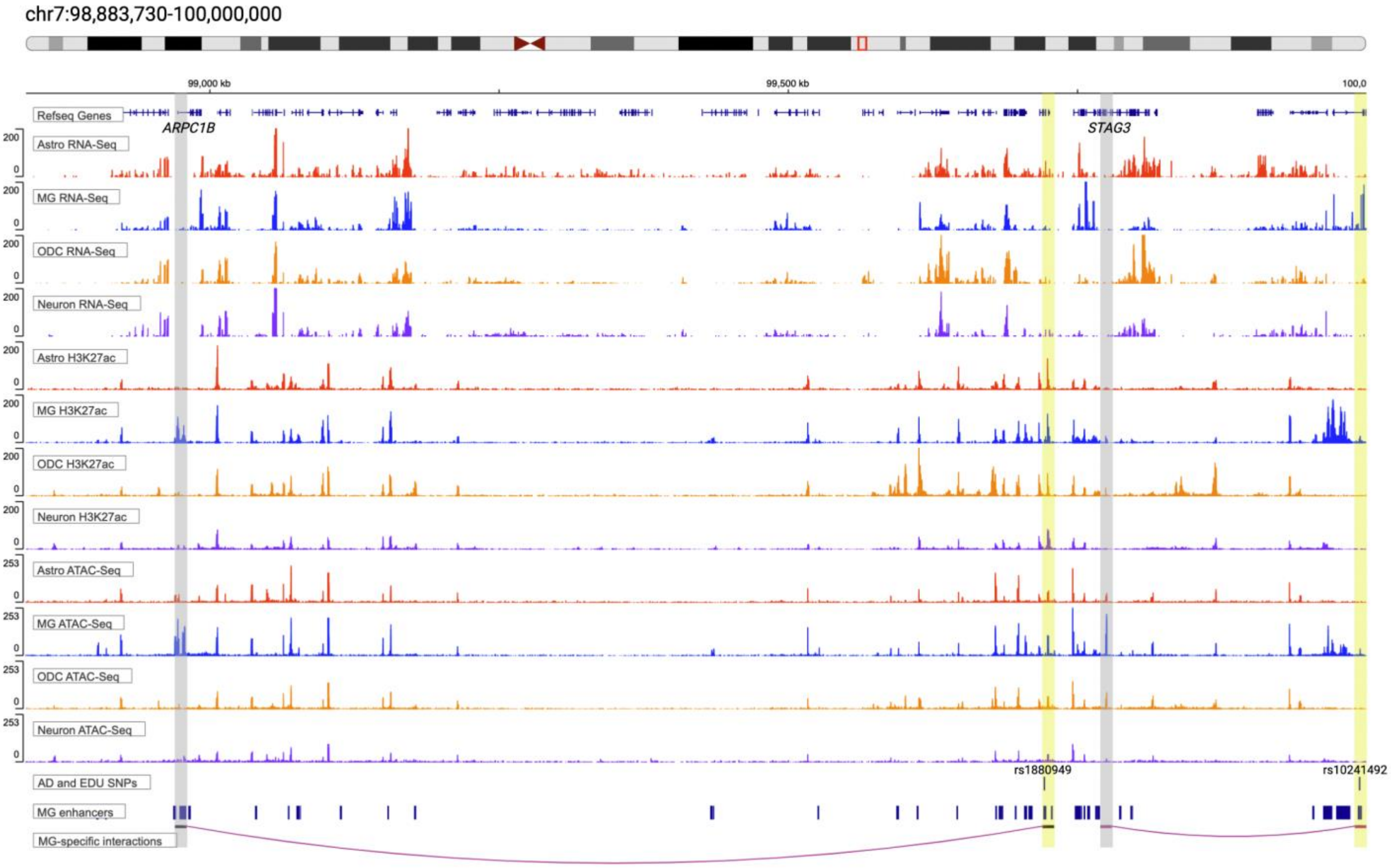


**Figure S1.** An illustrative example at the *APRC1B-STAG3* locus on chromosome 7 with a microglia-specific interaction. The top panel shows the gene track. The middle panels show RNA-seq, H3K27ac and ATAC-seq tracks for the four brain cell types. The bottom panels show AD and EDU GWAS SNPs, microglia enhancer regions, and two microglia-specific interactions identified by SnapHiC-G but not SnapHiC. One microglia-specific interaction links the promoter region of *ARPC1B* (highlighted in grey) with an AD-associated SNP rs1880949 (highlighted in yellow); the other one links the promoter region of *STAG3* (highlighted in grey) with an EDU-association SNP rs10241492 (highlighted in yellow). Both genes highlighted showed microglia-specific gene expression. The anchors of these interactions also showed stronger H3K27ac ChIP-seq and ATAC-seq signals in microglia.

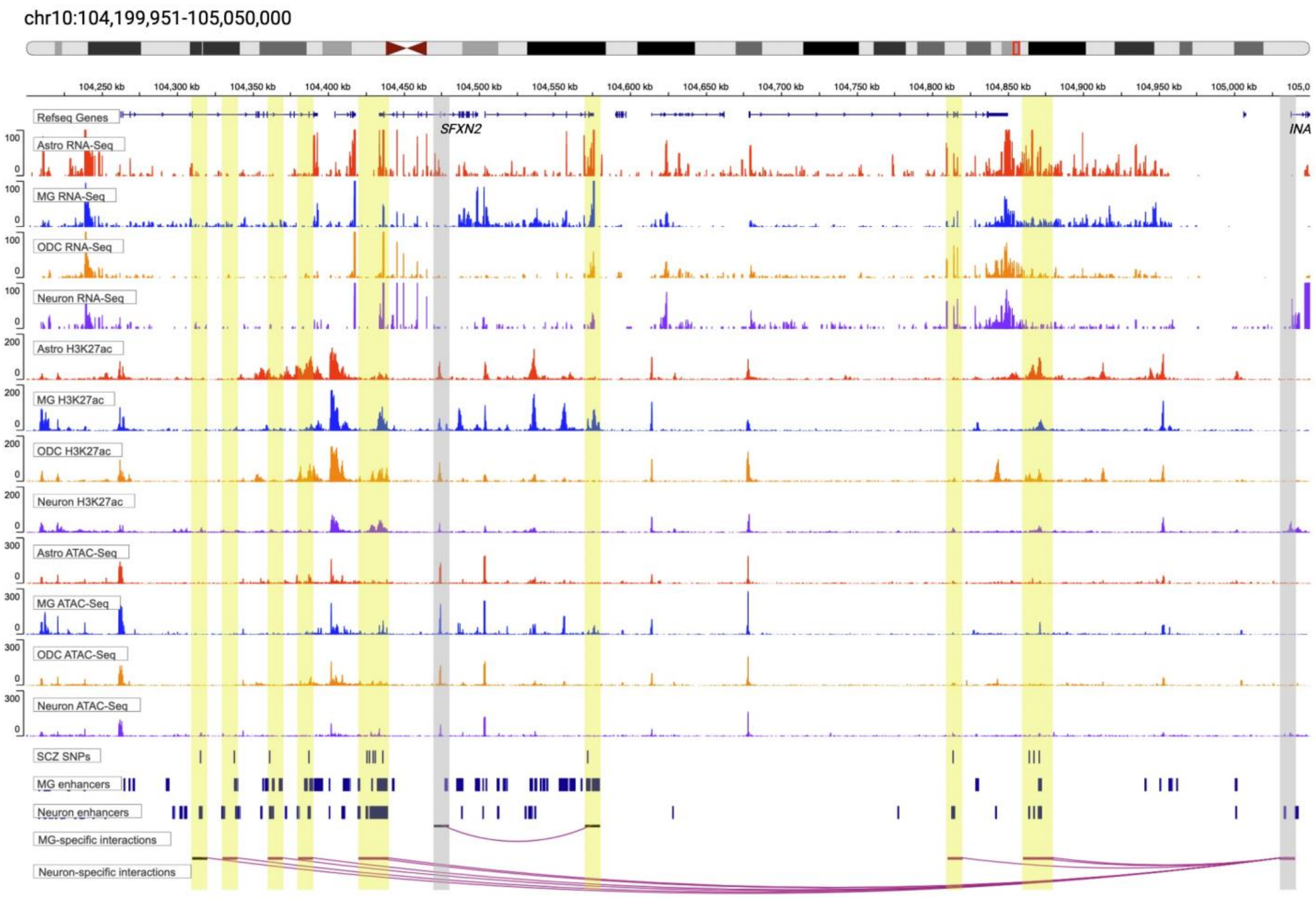


**Figure S2.** An illustrative example at the *SFXN2-INA* locus on chromosome 10 with neuron- and microglia-specific interactions. The top panel shows the gene track. The middle panels show RNA-Seq, H3K27ac and ATAC-seq tracks for the four brain cell types. The bottom panels show SCZ GWAS SNPs, enhancer regions in microglia and neurons, and cell-type-specific interactions identified by SnapHiC-G but not SnapHiC. A microglia-specific interaction links the promoter region of *SFXN2* (highlighted in grey) with a SCZ-associated SNP (highlighted in yellow); multiple neuron-specific interactions link the promoter region of *INA* (highlighted in grey) to SCZ-association loci (highlighted in yellow). Both genes highlighted showed cell-type-specific gene expression in corresponding cell types. The anchors of these interactions also showed stronger H3K27ac ChIP-seq and ATAC-seq signals in matched cell types.

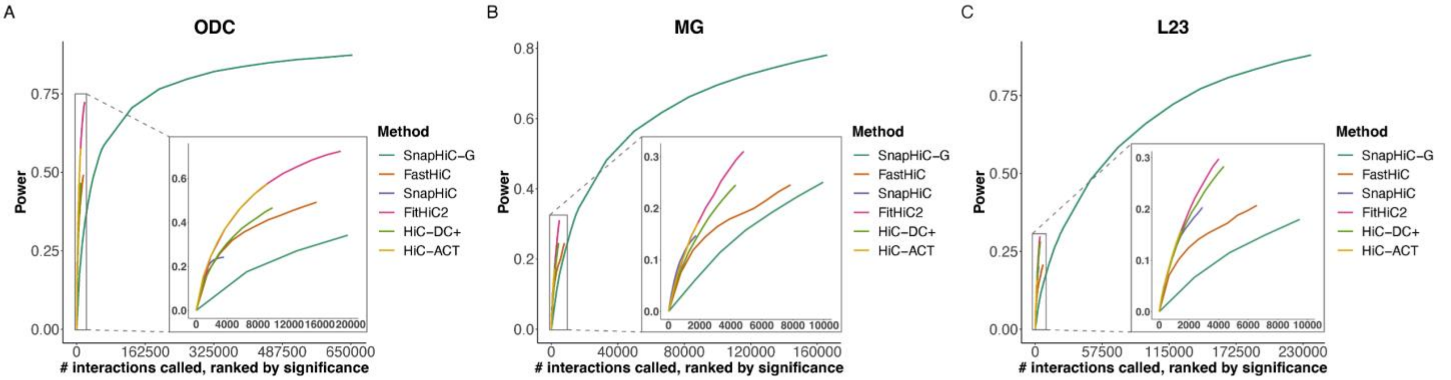


**Figure S3.** Power curves with (a) oligodendrocytes, (b) microglia, and (c) L2/3 neurons. Interactions were ranked by significance on the x-axis and power was evaluated with the corresponding number of top interactions. The lower right hand corner smaller figures are zoomed-in views of the top 20,000 interactions from oligodendrocytes and top 10,000 interactions from microglia and L2/3 neurons.

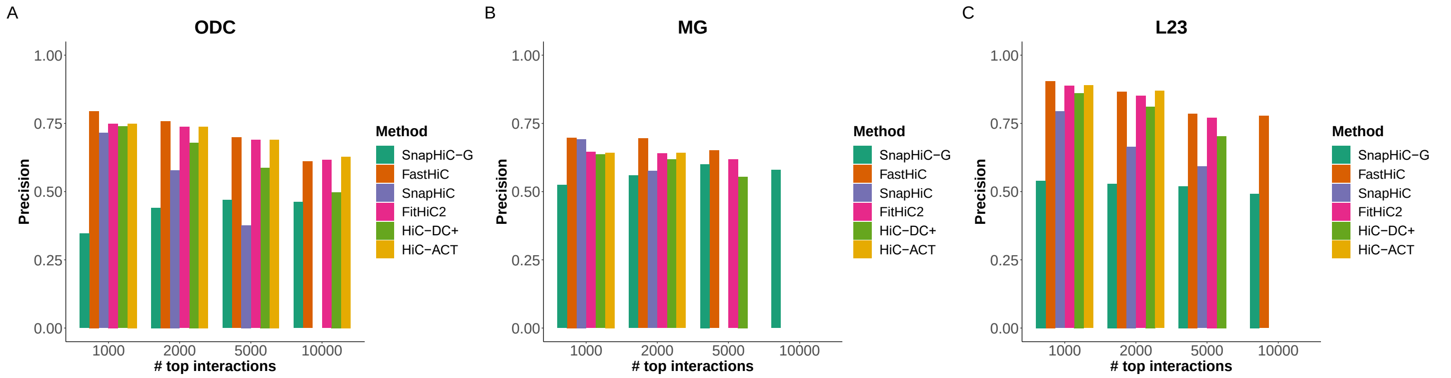


**Figure S4.** Precision barplots for three brain cell types. Precision bar plots with (a) oligodendrocytes, (b) microglia, and (c) L2/3 neurons. Results shown were for the top 1,000, 2,000, 5,000, and 10,000 interactions ranked by significance. Some bars are missing because the number exceeds the number of interactions called by that method.

**Table S1. Overlapped interactions in human brain cell types.**

A. Four-by-four table of two-way overlapped interactions.

| | Astro | MG | ODC | L23 |
|---|---|---|---|---|
| **Astro** | | 78,203 | 93,709 | 100,585 |
| **MG** | 80,727 | | 83,345 | 89,893 |
| **ODC** | 98,805 | 84,783 | | 105,034 |
| **L23** | 126,071 | 106,096 | 122,473 | |

Element (*i, j*) (e.g. (1, 3)) is the number of the interactions identified in the *i*-th (e.g., astrocytes) cell type that overlap with the interactions identified in the *j*-th (e.g., oligodendrocytes) cell type.

B. Four-by-seven table of three-way overlapped interactions.

| | MG ODC | MG L23 | ODC L23 | Astro ODC | Astro L23 | Astro MG |
|---|---|---|---|---|---|---|
| **Astro** | 63,681 | 64,752 | 76,188 | | | |
| **MG** | | | 64,345 | 63,290 | 65,081 | |
| **ODC** | | 66,665 | | | 78,998 | 66,410 |
| **L23** | 78,135 | | | 92,437 | | 79,696 |

Element (*i*, *j*) (e.g., (1, 3)) is the number of the interactions identified in the *i*-th (e.g., astrocytes) cell type that overlaps with the interactions identified in both cell types *j* (e.g., oligodendrocytes and L2/3 neurons).

C. Number of cell-type specific interactions.

| Astro | MG | ODC | L23 |
|---|---|---|---|
| 14,440 | 39,220 | 23,853 | 65,819 |

Column *i* is the number of interactions that are identified in the *i*-th cell type that does not overlap with interactions in any of the other cell types.

**Table S2. Running time and memory for SnapHiC-G step C and step D.**

| Genome | # cells | Bin pair distance | Cell type | MaxRSS/node (Gb) | runtime per CPU (hour) | # CPUs | # tasks/node | # Nodes | Memory-per-task |
|---|---|---|---|---|---|---|---|---|---|
| mm10 | 500 | 20Kb-1Mb | mESC | 12.72 | 0.59 | 100 | 25 | 4 | 0.51 |
| mm10 | 742 | 20Kb-1Mb | mESC | 9.00 | 0.78 | 150 | 10 | 15 | 0.90 |
| hg19 | 338 | 20Kb-1Mb | Astro | 11.19 | 0.57 | 60 | 20 | 3 | 0.56 |
| hg19 | 323 | 20Kb-1Mb | MG | 7.61 | 0.68 | 60 | 10 | 6 | 0.76 |
| hg19 | 1038 | 20Kb-1Mb | ODC | 10.84 | 1.27 | 80 | 10 | 8 | 1.08 |
| hg19 | 261 | 20Kb-1Mb | ODC | 7.88 | 1.77 | 60 | 10 | 6 | 0.79 |

All scHi-C data are at 10 Kb resolution.
